# Neural signatures for temporal order memory in the macaque medial posterior parietal cortex

**DOI:** 10.1101/2023.08.17.553665

**Authors:** Shuzhen Zuo, Chenyu Wang, Lei Wang, Zhiyong Jin, Xufeng Zhou, Ning Su, Jianhua Liu, Thomas J. McHugh, Makoto Kusunoki, Sze Chai Kwok

## Abstract

Episodic memory involves encoding and remembering the order of events experienced over time. Previous work examining the mechanisms of temporal-order memories has focused on the hippocampus and prefrontal cortices but has largely ignored the memory ensembles in the medial posterior parietal cortex (mPPC). Combining *in vivo* multi-unit electrophysiology and a temporal-order judgment task with cinematic material in the macaque, we find that mPPC neuronal activity reflects temporal context both during encoding and recall. During learning, mPPC neuronal ensembles encode the temporal information as well as contextual information embedded in the videos, whereas at retrieval these neurons fire in synchrony prior to memory-guided decisions. Moreover, the similarity between encoding and retrieval correlates with the animal’s performance. A control experiment further ruled out eye saccades, fixation, and scan path for their confounding roles in the neural results. Together, these data suggest that neurons in mPPC track the passage of time and contextual changes, thereby orchestrating for successful temporal-order memories at retrieval.

**SIGNIFICANCE:** Remembering the temporal order of events is a vital function of the brain. This study provides the first evidence for the retrieval of temporal order of episodes at both single neuron and ensemble levels in the primate parietal cortex. The study’s key breakthrough is the revelation that ensembles of time perceptive neurons, acting as “temporal context cells”, can encode a spectrum of time constants and contexts for the experienced past and subsequently utilize this temporal record to discriminate the chronological order of past experiences. These results enhance theoretical frameworks for how encoded temporal past is maintained and reconstructed during memory retrieval.

## INTRODUCTION

Remembering the temporal order of experiences is a fundamental feature of episodic memory ^1–4^. Ample research has implicated the MTL in different aspects of temporal memory, with evidence from a large number of studies suggesting that neural activities in the MTL, especially the hippocampus, mediate the formation of temporal order memories ^5–10^, potentially via sequence generation ^11^. While temporal information about past events may be present within the hippocampus ^12–14^, data also suggests that the computation for temporal memory and order of events may also involve the lateral PFC ^15,16^ and/or the posterior parietal cortex ^17,18^. Given the precuneus’ role in temporal context ^19^, temporal memory ^20^ as well as in forming new memories ^21,22^, the medial posterior parietal cortex (mPPC) may be expected to serve as a plausible neural hub in the computation for the retrospective remembering of temporal order of experiences.

Theorists posit that successful memory of temporal order necessitates the brain linking temporal contextual information across memory phases. According to the temporal context model ^23–25^, when multiple events occur close together in time, they generate multiple sequences across the overall population which may overlap with each other depending on the temporal distance between these events. These sequences provide a readout of recent events and their temporal history. Empirical evidence on time cells and temporal context cells that can encode the passage of time and organize incoming information ^26,27^ corroborate the predictions of these theories for the encoding phase of the experiences. However, while the temporal coding properties of “time cells” offer a suitable mechanism by which time may be encoded, it is not clear whether and how such temporal context signals ^26^ established at encoding would later be reactivated and utilized to help determine the temporal order for different episodes of experience at retrieval.

To address these questions, we recorded multi-neuron activities from the monkey mPPC (the precuneus) while the animals made temporal order judgment of two still frames extracted from encoded videos. We found that neurons in the mPPC encode the passage of time and engender a spectrum of time contextual constants during encoding (videos) and similar pattens of activity are reinstated during the subsequent recollection of memories, with synchronous neuronal activity reflecting the accumulation of mnemonic information for higher memory accuracy. Moreover, we ascertained that oculomotor behavior cannot explain away neuronal activities related to temporal order memory. We also found that the putative temporal context cells are context-dependent depending on the encoded material.

## RESULTS

In total, we completed 42 daily sessions of data collection from Monkey Jupiter, 21 sessions from Monkey Mercury (Main experiment), and 21 sessions from Monkey Mars (Experiment 2). We recorded 874 neurons in total from the three animals. In the main experiment, we recorded 676 single neurons from 2 of the macaque monkeys (401 cells from Jupiter and 275 cells from Mercury). In experiment 2, we acquired 198 neurons from Mars. On average, 9.548±2.276, 13.095±4.265, and 9.429±2.014 neurons per session were recorded simultaneously for Jupiter, Mercury, and Mars respectively.

### Forward memory search during temporal order judgment of cinematic events

In the main experiment, after watching an 8-s naturalistic video, monkeys performed a temporal order judgment task during which they were required to judge the earlier frame among two frames extracted from the video. We controlled temporal similarity for the two conditions, thus having two delayed conditions here: immediate TOJ for frames taken from first half of the video (5^th^ and 90^th^ frames) vs. delayed TOJ for frames taken from latter half of the video (95^th^ and 180^th^ frames), Figure 1A. The performance of both monkeys was significantly higher than the chance level and remained steady through the experiment for the two conditions (Jupiter: immediate = 66.6%, delayed = 69.8%; overall mean = 67.7%, against chance *P* < 0.001, Mercury: immediate = 62.7%, delayed = 69.8%, overall mean = 64.2%, against chance *P* < 0.001), Figure 1C. We then compared the RTs under different chosen frame locations and replicated our previous results ^28^, showing that RTs were faster for frames chosen from the early part of the video (one-way ANOVA, Jupiter: F(3, 10171) = 52.91, *P* < 0.001, post hoc test, 5^th^ frame vs 90^th^ frame, *P* < 0.001, 95^th^ frame vs 180^th^ frame, *P* < 0.001; Mercury: F(3, 3195) = 5.86, *P* < 0.001, post hoc test, 5^th^ frame vs 90^th^ frame, *P* = 0.006, 95^th^ frame vs 180^th^ frame, *P* = 0.041) and a numeral trend of increased RTs as a function of the four chosen frame locations (Figure 1D). In Experiment 2, Mars also showed good memory performance (immediate = 79.07%, delayed = 77.35%; overall mean = 78.21%, against chance *P* < 0.001) and did not show any reaction time effect for across context TOJ responses, consistent with our previous report ^28^.

**Figure 1.**
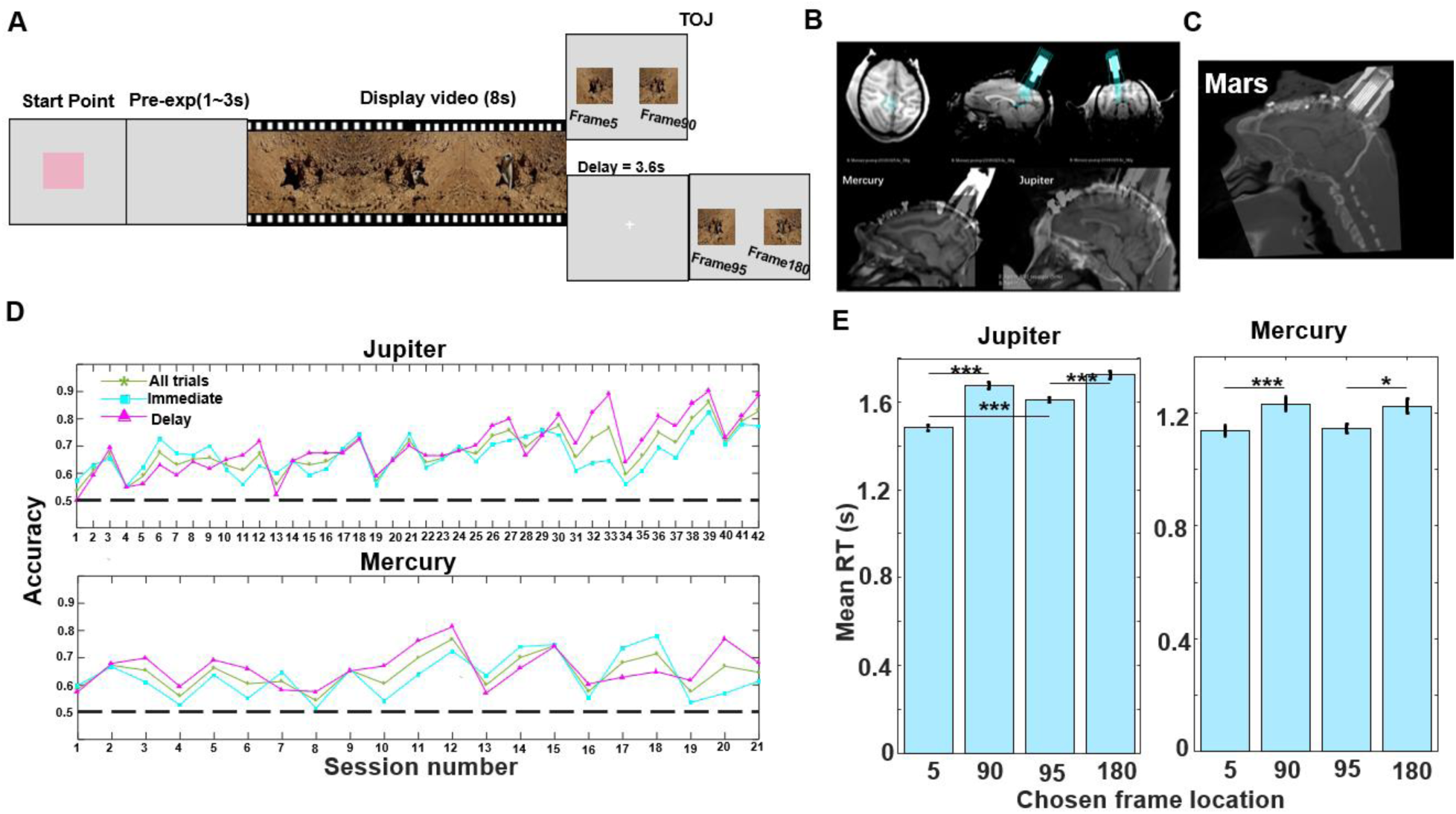
Paradigm, recording system and sites, and behavioral results. (A) Temporal order judgment task. In each trial, the monkey watched an 8-s video, and following a 3.6-s delay or 0-s delay, the monkey judged which frame they had seen earlier to get a reward. (B) Upper panel: Anatomical MRI images showing the position of recording chamber (in blue) implanted in the medial posterior parietal cortex. Bottom panel: CT images visualizing the location of the electrodes when aligned with MRI images for monkey Mercury and Jupiter (Main experiment) (C) A CT image visualizing the location of the electrodes of Mars in alignment with T1 MRI image (Experiment 2). (D) Accuracy percentage across all sessions for all trials (green star), immediate condition (cyan square) and delayed condition (magenta triangle) of Jupiter and Mercury. (E) Mean reaction times of different chosen frame locations. *** *P* < 0.001, * *P* < 0.05.

### Identification of temporal context cells during encoding period

Firstly, we asked if mPPC neurons are involved in the temporal memory encoding. We set out to find temporal context cells that help code for time and build a temporal contextual record of the past ^26^. To examine whether the mPPC neurons are involved in the representation of temporal context information (cf. temporal context cells in monkey entorhinal cortex ^26,29^), we fitted the spike data during the encoding period (videos) with three different models for each neuron (see ‘Temporal context cell fitting’ in STAR Methods). We first fit the firing pattern of each cell with a convolution of Gaussian and an exponential function ^26^. There are two key parameters that would determine the shape of distribution. One parameter is the mean of Gaussian *µ*, which indicates the response latency of each neuron. Another parameter is the time constant of the exponential function *τ*, which represents the time each neuron needs to relax back to 63% of the peak firing rate (relaxation time). For comparison, we also fitted our data with the constant model and the Gaussian model. Using the likelihood ratio test for model comparison, we identified 111 neurons (out of 676) as temporal context cells. Among these temporal context cells, 63 neurons increase their firing rate at the beginning of video (see an example neuron in Figure 2A, left), while 48 neurons decrease their firing rate (Figure 2A, right). The normalized firing rate pattern for the whole population of temporal context cells shows that most neurons in the mPPC reacted to the video material within a short period of time (Figure 2B). When we looked at the relationship between the cells’ response latency and their relaxation time, a majority of the neurons respond to the video onset within 1 s and decrease their firing rate with different latency ranging from 0 s to a hypothetical 20 s (Figure 2C), highly resembling the patterns previously reported for entorhinal temporal context cells ^26^. It is possible that the temporal context cells as a whole cover a broad spectrum of time, representing the temporal contextual information embedded in the videos.

**Figure 2.**
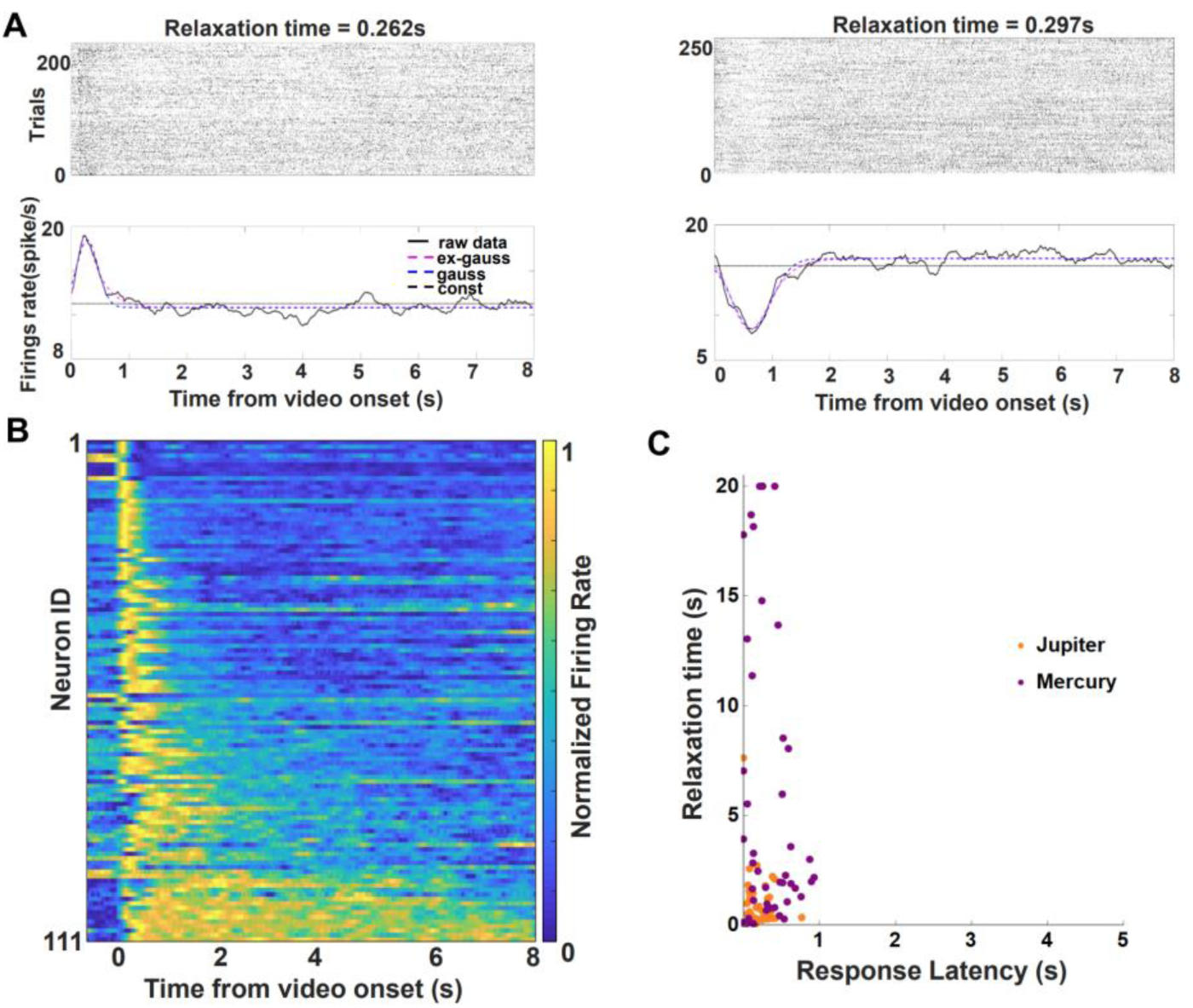
Temporal context cells during encoding (video) period. (A) Two examples of temporal context cells during video-watching period. Spike raster (upper) and peristimulus time histograms (PSTHs, lower) plotted against the onset time of video- watching. In the PSTHs, the solid black line indicates the smoothed firing rate. The dashed pink line, the dashed blue line, and the dashed black line indicate the fitting result of the ex- Gaussian, gaussian and constant model, respectively. Relaxation time means the time interval between peak firing rate and when the neurons returned 63% of the way to baseline. The example on the left increases its firing rate at the beginning of video, whereas the example on the right decreases its firing rate. (B) A heatmap of normalized firing rate of 111 temporal context cells, relative to video onset, sorted by their relaxation time. Some neurons relaxed back to baseline slowly, covering the whole duration of the video. Overall, the neurons relaxed back to baseline with a broad spectrum of decay rates. (C) The plot of each temporal context cell’s response latency and relaxation time. The response latency of all temporal context cells is less than 1s, while the relaxation time ranges from 0 to 20 s. The dots indicate the temporal context cells recorded from monkey Jupiter (66 neurons, orange) and monkey Mercury (45 neurons, purple), respectively.

### Cell ensembles carry information about passage of time

We next assessed how these neurons represent the passage of time during encoding as a population. We trained a linear discriminant analysis (LDA) decoder to estimate the time following the presentation of a video. To the extent the predicted time bin for out-of-sample data is close to the actual time bin, we inferred that the population response carried information about time. In each trial, we took 250 ms as the time bin and divided each 8 s video into 32 bins. We trained the classifier with odd trials and tested the temporal information in even trials. The LDA results show that the mean absolute decoding errors (Jupiter: 1.58 s; Mercury: 1.60 s) were reliably lower than the decoding error after 1000 permutations (Jupiter: mean = 2.661 s, SD = 0.183 s, z score = -5.907; Mercury: mean = 2.664 s, SD = 0.199 s, z score = -5.347, Figure 3A-B, 3D-E). When we looked at the decoding error in each time bin, the decoding error of the first 1s time bins was much lower than the rest of the time bins for both monkeys (Figure 3C, 3F, 8H). In addition, we replicated these results on a third monkey in experiment 2 (Mars: absolute decoding error = 1.87 s, which was lower than permutated mean = 2.663 s, SD = 0.049 s, z score = -16.184, Figure 8F-G). It is noteworthy that in this 2-clip video design, the posterior probability increased again around the onset of clip 2 (at 4^th^ second), indicating that the cell ensembles contain information for both the passage of time as well as the change in context. To ensure that the decoding results are not contaminated by the initial period of video, we repeated the LDA analysis but now cut the time bins from the start of encoding at the step of 0.5 s. This control analysis shows that the absolute decoding error was still significantly lower than permutation errors in all three monkeys (Figure S2), implying that the mPPC neurons carry information about the temporal unfolding of the videos.

**Figure 3.**
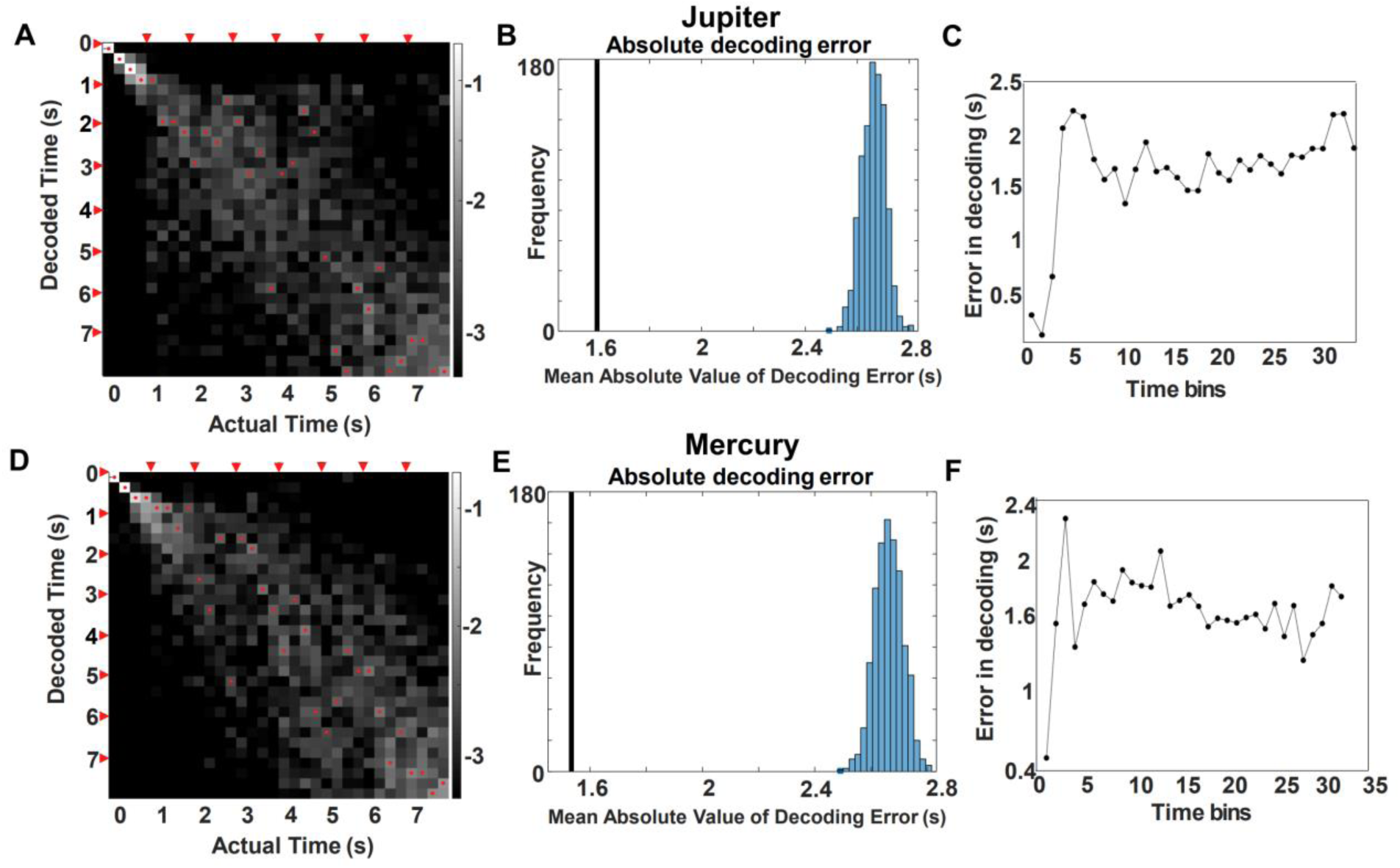
mPPC neurons encode passage of temporal information in videos. (A) Decoder performance based on the entire population of mPPC neurons in monkey Jupiter. The x-axis indicates the actual time bin, and the y-axis indicates the decoded time. Each time bin represents 250 ms of the video, and each 8-s video was divided into 32 bins. The grayscale indicates the posterior probability in the logarithmic scale, with a lighter shade indicating higher probability. The decoded time with the highest probability in each time bin is marked with a red dot. Each second was marked with red arrows along the axis. (B) The actual decoded errors are shown by a vertical line (1.58 s for Jupiter, 1.6 s for Mercury) and the distribution of errors obtained from 1000 permutations. In both monkeys, actual errors are significantly smaller than the errors by permutation, indicating a successful decoding. (C) The absolute decoding error in each time bin. For both monkeys, the error was relatively lower at the beginning of the video (around first 0.5 s; see also Figure S2). (D) – (F) same as (A-C) for Mercury.

### mPPC neurons mediate TOJ processes

We then verified whether mPPC neurons code for memory during temporal order of judgement. Given the region’s role in memory recollection ^30^, we expected to observe neuronal signals to be related to memory processes and performance. To address this question, we ran a Poisson GLM taking into account the within-session block number, reaction time, response outcome (correct vs incorrect), conditions (immediate vs delayed condition), touch side (left vs right), and events within a trial including preparation, encoding period, delay period, TOJ period, feedback (reward vs blank), and ITI period. We identified the neurons that are significantly responsive to the TOJ event, which we refer to as “TOJ cells” (n = 447: Jupiter: 278 cells, Mercury: 169 cells). Together with the non-TOJ neurons (n = 229: Jupiter: 123 cells, Mercury: 106 cells), we fisher transformed their GLM coefficients (adjusted *R^2^* values) into normal distribution and confirmed that the *R* values are significantly larger for the TOJ cells than for non-TOJ cells in the main experiment (Jupiter: two-sample t-tests, t (399) = 7.806, P<0.001, Cohen’s d=0.793; Mercury: t (273) =16.640, *P*<0.001, Cohen’s d=2.029) (Figure 4A). In the control experiment in which we utilized 2-clip videos for encoding, we found the proportion of TOJ cells was lower for this third monkey (see Monkey Mars in Table S1and in Figure 8E: TOJ cells, n = 58, non-TOJ cells, n = 140). This discrepancy is in line with our previous findings that macaques apply two different memory decision strategies for within-context (1-clip) vs. across-context (2-clip) encoded material in terms of their respective rates of rise of information accumulation^28^.

**Figure 4.**
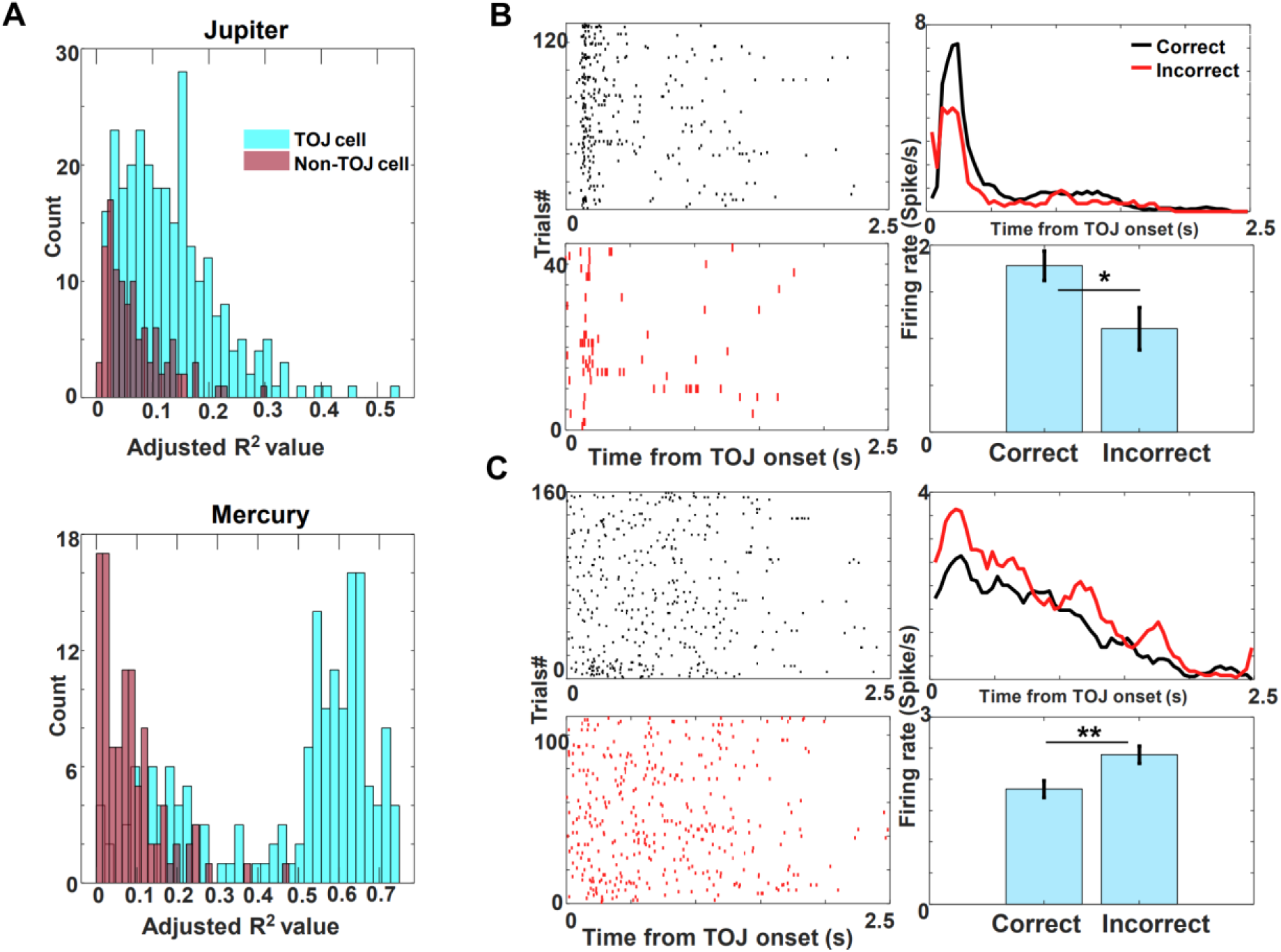
TOJ cells and neuronal activities reflecting memory correctness for temporal order judgment. (A) Histogram of R^2^ values from Poisson GLM for TOJ cell (cyan) and non-TOJ cell (brown) of two monkeys. 278 TOJ cells and 123 non-TOJ cells were identified for Jupiter, and 169 and 106 cells identified for Mercury, respectively. (B) An example neuron that showed higher firing rates for correct trials than incorrect trials (one- tailed t-test, P = 0.027). Left panels show the raster display of the neuron’s spike activity in correct trials (top) and incorrect trials (bottom). Top right panel shows the mean response of the example neuron in both correct trials and incorrect trials. Mean firing rates in correct and incorrect trials are shown in black and red, respectively. Bottom right panel shows the group average (n = 36 cells). (C) Same as (B), but with an example neuron that showed higher firing rates for incorrect trials than correct trials (one-tailed t-test, P = 0.003). Bottom right panel shows the group average (n = 36 cells).

Moreover, when we examined the firing rate for correct trials versus incorrect trials during the TOJ period, 72 (∼10.7%) neurons differed in their firing rates between correct response and incorrect response (one-tailed t-test, all *P*s<0.05). There were two types of neurons: one type that increased its firing rate for correct decisions (n = 36, Figure 4B) and another one that increased its firing rate for incorrect decisions (n = 36, Figure 4C). This opposing response pattern has been previously reported in hippocampal neurons for repeat vs. novel still images ^31^.

### Spike train synchrony tracks evidence accumulation for TOJ decision

Given that mPPC neurons are involved in memory retrieval, we next examined their activity during the recall period. We took the temporal dynamics (relative spike timing) across the spike trains of multiple neurons into consideration to reveal how spike synchrony tracks the processes of temporal order judgment. We calculated real-time SPIKE-distance based on the temporal information of firing for all neurons recorded within a session ^32^, with lower SPIKE-distance values reflecting greater synchrony, to examine the degree of synchronous activity time-locked to either the TOJ onset (i.e., when two probe frames shown on screen) or the TOJ response offset (i.e., when the monkey made the decision). During the TOJ onset period, the synchrony level increased and reached its peak within 200 ms, indicating that mPPC neurons work in concert in a progressive manner after the probe frames initiated the retrieval process. This spike train synchrony asymptotes very quickly (< 200 ms), but we did not observe persistent differences in correct vs. incorrect responses during this initial period (Figure 5A-B top). In contrast, during the time just prior to the monkeys’ responses (TOJ offset), the synchrony level in correct trials is significantly, and persistently, higher than in incorrect trials as the monkey’s decision time approaches (Figure 5A-B bottom), indicating that the population synchrony is stronger during successful TOJ decisions. This is a continual process with statistical differences more densely observed 400-500 ms prior to manual responses made by the monkeys, implying the neurons fire in synchrony to regulate the accumulation of mnemonic information and result in higher memory accuracy. Additionally, we observed differences in the pattern of synchrony level during the TOJ period for both the immediate and delayed conditions. The synchrony level is consistently higher for the delayed condition compared to the immediate condition both time-locked to TOJ onset (Figure 5C-D top) or at monkeys’ responses (Figure 5C-D bottom). Since memory traces for trials without a retention delay ought to be stronger than those with a longer retention delay, these data imply that a high synchronization among spike trains is a neural proxy for memory strength.

**Figure 5.**
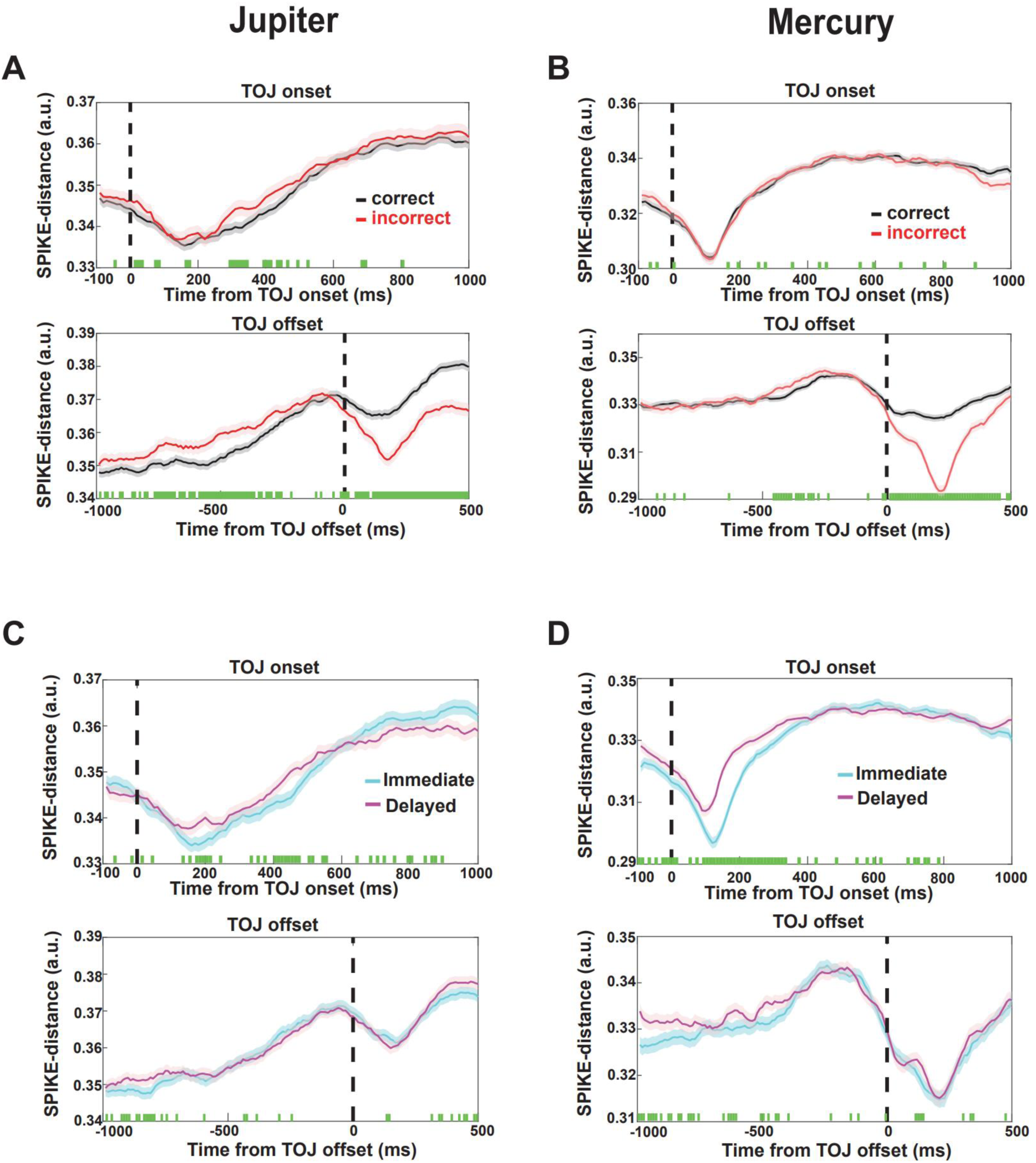
Multivariate SPIKE-profiles at TOJ tracks the process of memory retrieval and support forward search. (A) – (B) Upper panels show the fluctuation of spike trains synchrony during TOJ. We time-locked 0 ms to the onset of TOJ images. Lower panels show the dynamic of spike trains synchrony with zero time-locked to the moment when a TOJ decision was made (1000 ms before to 500 ms after a TOJ decision). Black lines show the mean value of SPIKE-distance in correct trials, while red lines represent those for incorrect trials. Green dots at the bottom of each panel indicate a significant difference between correct and incorrect trials for the 10-ms time bin (paired samples t-tests, P <0.01). We also noted a divergence in SPIKE-distance after the response, which might be attributed to post-decision, non-mnemonic processes. Shaded areas represent SEM across sessions. (C) – (D) We compared the spike trains synchrony for the two experimental conditions (immediate vs. delayed condition) for the two monkeys. The synchrony level is consistently higher for the immediate condition than for the delayed condition. Legends are the same as (A-B).

### Similarity of neural activity between encoding and retrieval reflects temporal memory judgement

Since encoding-retrieval neural similarity has been reported to modulate memory success ^33,34^, we used population-based similarity to quantify the link between the encoding and retrieval phases of the task. For each trial, we constructed N-dimensional population vectors for the firing rate of encoding and retrieval period respectively (N refers to the number of neurons recorded in that session). Mahalanobis distance was calculated between the population vector of encoding and retrieval periods as a measure of dissimilarity in the pattern of firing across the two stages. To compare the Mahalanobis distance across sessions with varying number of cells (ranging from 5 to 21 neurons per session), we divided the Mahalanobis distance by two times the number of cells in that session and obtained a Distance Index ^6^. As shown in several example trials, the Mahalanobis distance was consistently smaller during correct trials (Figure 6A). Indeed, two-samples *t*-tests revealed a significant lower Distance Index (that is, more similarity) for correct trials (Jupiter: mean = 0.517, SD = 0.726; Mercury: mean = 0.913, SD = 0.543) than incorrect trials (Jupiter: mean = 0.664, SD = 0.701; Mercury: mean = 1.086, SD = 0.821) (Jupiter: one-tailed *t*_10405_ = -9.722, *P* < 0.001, Cohen’s d = -0.206, 95% CI: -Inf to -0.122; Mercury: one-tailed *t* _3605_ = -7.589, *P* < 0.001, Cohen’s d = 0.249, 95% CI: -Inf to -0.135; Figure 6B). Our finding of spike pattern reinstatement during TOJ aligns with data from single neurons in human patients that suggest the population states observed during initial experience are reinstated during subsequent recall ^35–37^. Overall, we showed that mPPC neurons carry memory related information and their engagement in encoding-retrieval neural similarity supports successful memory for temporal order of video events.

**Figure 6.**
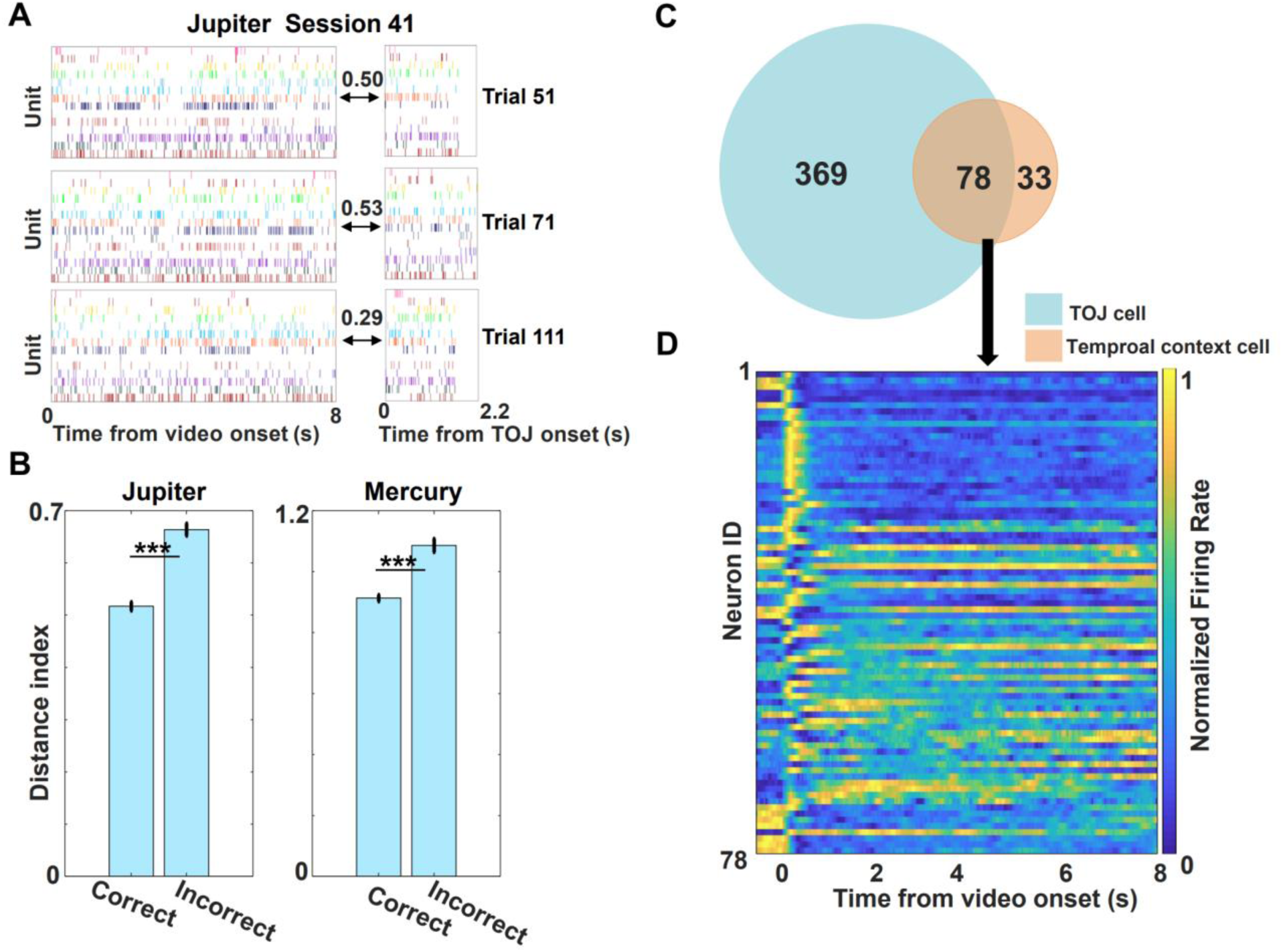
Population based Mahalanobis distance and TOJ cells ∩ temporal context cells overlap. (A) Examples of pattern changes across encoding and retrieval within one session. The data in these panels is taken from trial 51, trial 71, and trial 111 of session 41 of Jupiter illustrating the population activity from encoding and TOJ period. The numbers shown are the Distance Index between the two periods for each trial. (B) Similarity of ensemble responses for correct and incorrect trials. Incorrect trials show a higher Distance Index value than correct trials, indicating a higher pattern similarity between encoding and retrieval is crucial for successful memory recall. * *P* <0.05, ** *P* <0.01, *** *P* <0.001. (C) Venn diagram showing the cells (78 cells) that overlapped between temporal context cells (111 cells) and TOJ cells (447 cells). (D) A heat plot of the normalized firing rate of 78 cells overlapped between the temporal context cells and the TOJ cells.

Given that the temporal context neurons were found to be involved in the encoding of temporal context, these cells would have different relaxation times and be relaxing to the baseline at different rates to form a whole course temporal memory of the videos ^26^ via Laplace transform ^27^ (see Figure 2). We then examined the relationship between the temporal context cells and their functional relevancy with memory recollection of temporal order. We identified the TOJ cells that are also classified as temporal context cells, observing that ∼70.3% (78 out of 111 cells) of the temporal context cells belonged to the TOJ cells class (Figure 6C-D). A chi-square test revealed that this proportion is statistically significantly larger than what would be expected by chance (χ^2^(1) = 91.203, *P* < 10^-5^). Therefore, we inferred that the neurons’ ability forming a broad spectrum of time would also allow them to utilize prior temporal context at encoding to support the subsequent TOJ.

### Eye movement confounding effects regressed out from TOJ neuronal activity

To control for potential effects of eye movement during TOJ period, we trained a third monkey on this temporal order judgment task and recorded the monkey’s eye position, scan paths, and neuronal activities during the task simultaneously (see Movie S1). Knowing where the animal looked and scanned across the TOJ frames, we tracked neurons’ activities as a function of the monkey’s eye fixations and scan paths. The neurons firing rates changed during the time course during memory decision processes. We presented the firing rate as a function of either fixation or scan path for six example cells, separately for correct (Figure 7A) and incorrect responses (Figure 7B). From these, we can see that the animal overtly attended to the two still frames while they performed the temporal order memory judgement. We then ran a control analysis in which we included frequency of saccade, frequency of fixation, total scan path length, and total duration of fixation into a Poisson GLM model. The result showed that cells that included TOJ period as a significant variable changed from 58 cells to 41 cells (Table S1). Importantly, among these 41 TOJ cells, a considerable proportion of 70.1% (29 out of 41 cells) did not include any eye movement related parameters (note: 6 cells included fixation duration as a significant factor, 5 cells included frequency of fixation as a significant factor and only one cell included frequency of saccade). By comparison, when we examined the proportion of non-TOJ cells including eye movement parameters as a significant variable, a comparable 82.8% (130 out of 157 cells) did not include any eye movement related parameters in the final model (note: frequency of fixation: 21 cells, frequency of saccade: 4 cells, and fixation duration: 2 cells). Those results indicate that eye movement activities did not account for the neuronal activity for TOJ memory processes much differently from non-TOJ processes. These regression analysis results served to control for confounding roles of saccades, fixations, and scan paths and provided good validation that eye movement parameters *per se* did not account for our present TOJ related neuronal activities in an idiosyncratic manner.

**Figure 7.**
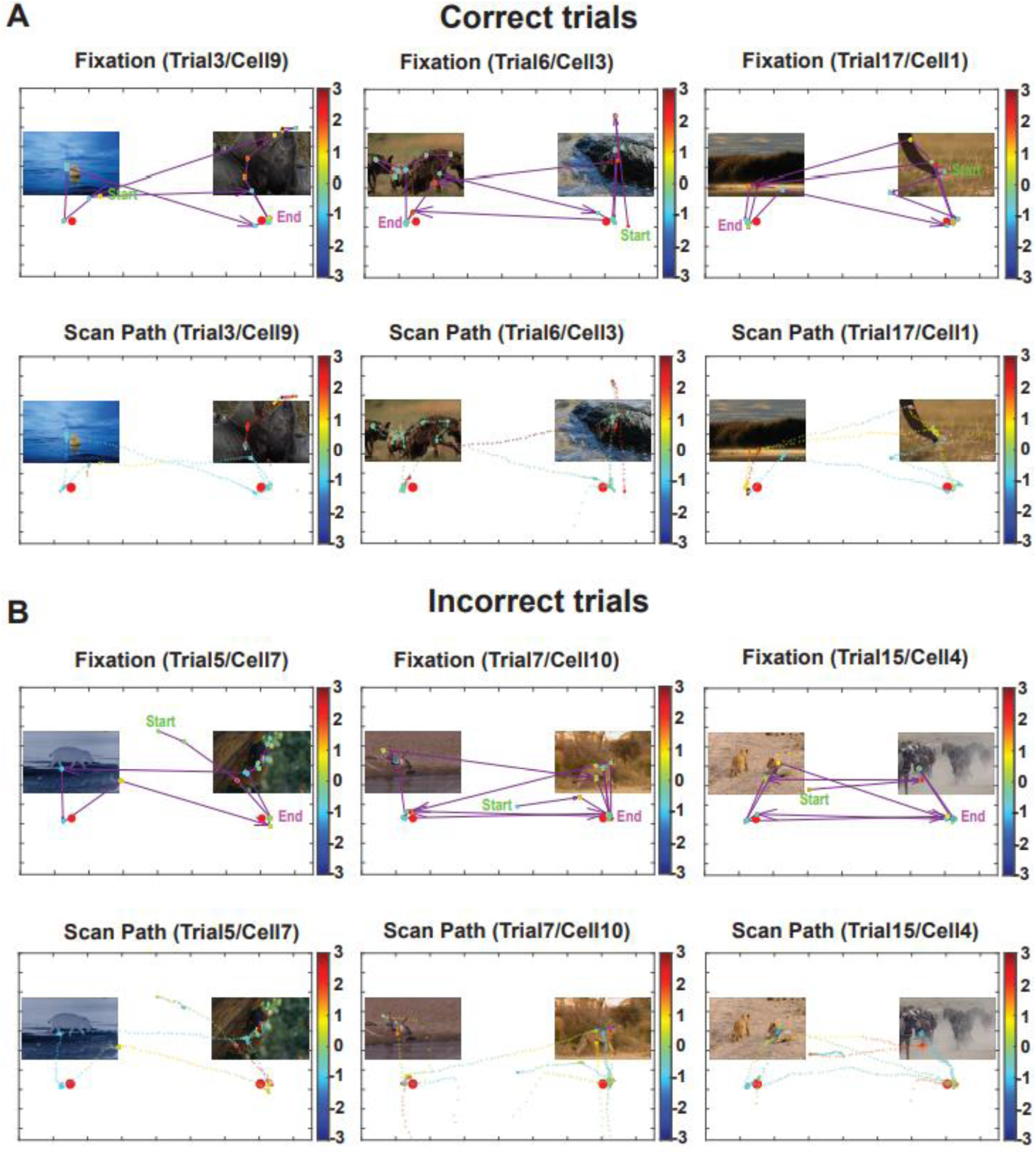
Eye position and corresponding firing rate. (A-B) Normalized firing rates of six example cells are plotted as a function of eye gaze fixation and saccadic scan path for three correct trials (A) and three incorrect trials (B). The normalized firing rate of the cells are shown with the colored discs in the fixation plots (upper rows) and with the colored dotted paths in the scan path plots (bottom rows). The color bars depict normalized firing rates. In the fixation plots, the beginning and conclusion of a trial are marked with “Start” and “End”; the arrows show the direction and length of saccades. See also Movie S1.

**Figure 8.**
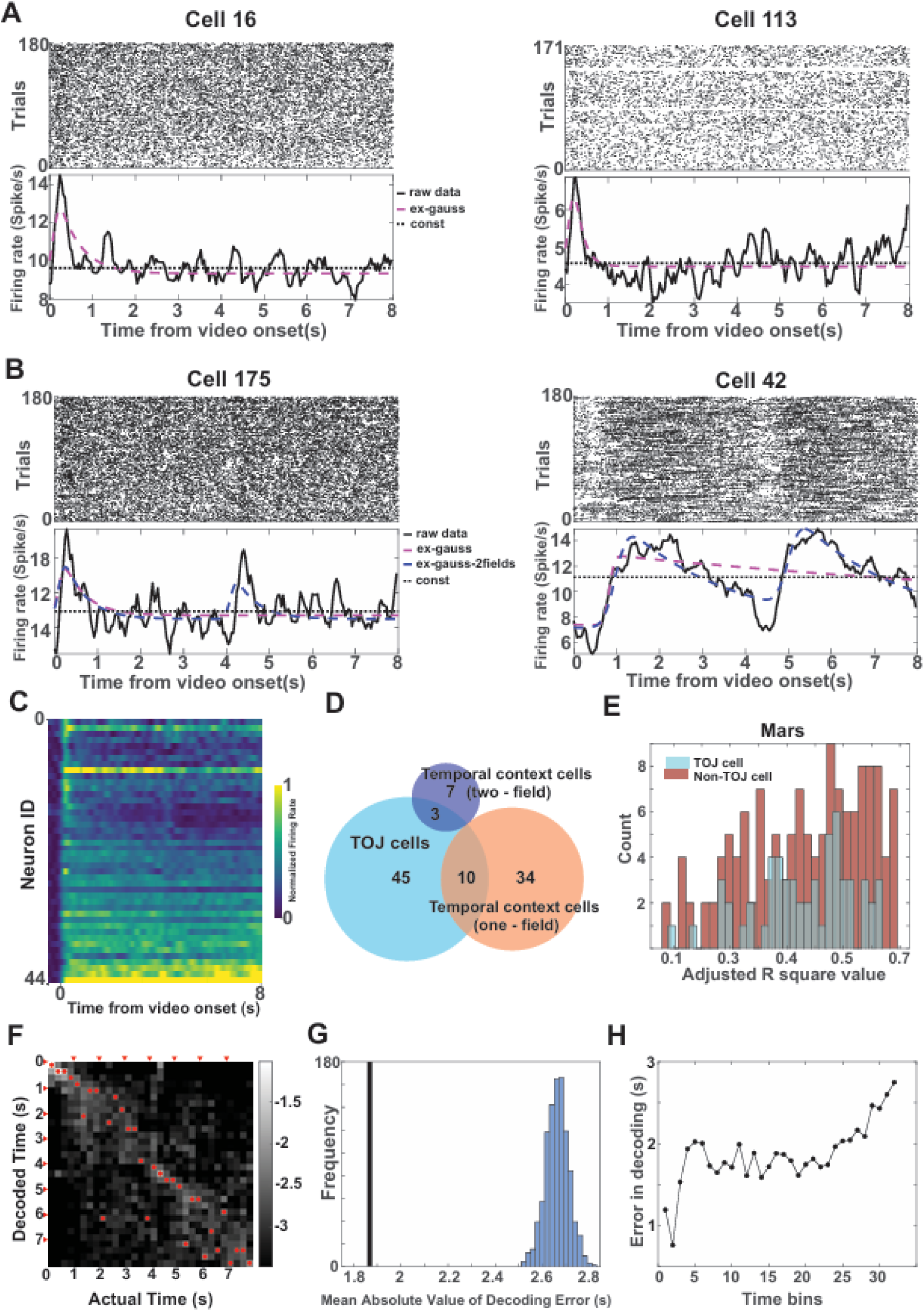
Summary result from experiment 2 showing temporal context cells, TOJ cells, and linear discriminant analysis (LDA) decoder. (A-B) Examples of one-field temporal context cells (A) and two-field temporal context cells (B). Spike raster (upper) and peristimulus time histograms (PSTHs, lower) plotted against the onset time of videos. The examples in (A) show only one temporal receptive field, which is aligned to the onset of the videos (hence, one-field cells) whereas the examples in (B) changed their firing in response to the onset of each clip within videos (two-field cells). In the PSTHs, the dashed red line, the dashed blue line, and the dashed black line indicate the fitting result of the ex-Gaussian, the two-field ex-Gaussian, and the constant model, respectively. The solid black line indicates the smoothed firing rate. (C) Heatmap of normalized firing rate of 44 one-field temporal context cells, relative to video onset. The neurons relaxed back to baseline with a broad spectrum of decay rates. (D) Venn diagram showing the numbers of temporal context cells (44 one-field cells and 10 two- field cells) and TOJ cells (n = 58), and their overlap. (E) Histogram of R^2^ values from Poisson GLM for TOJ cells (cyan, n = 58) and non-TOJ cells (brown, n = 140) of Mars. (F) Decoder performance based on the entire population of neurons in Monkey Mars. The x-axis indicates the actual time bin, and the y-axis indicates the decoded time. The grayscale indicates the posterior probability in the logarithmic scale, with a lighter shade indicating higher probability. The decoded time with the highest probability in each time bin is marked with a red dot. (G) The actual decoded errors are shown by a vertical line (1.87 s for Mars) and the distribution of errors obtained from 1000 permutations. Actual errors are significantly smaller than the errors by permutation, indicating a successful decoding. (H) The absolute decoding error value in each time bin. Consistent with Jupiter and Mercury, the error was relatively lower at the beginning of the video (around first 0.5 s; see also Figure S2). (I)

### Confirmation of temporal context cells being context-dependent

Finally, we interrogated the property of the temporal context cells using the data from experiment 2. In this experiment, we had the monkey perform the temporal order judgment task after encoding across-context videos. We hypothesized that the putative temporal context cells might show context-dependent coding patterns. Indeed, we found that while some mPPC neurons only had one temporal receptive field aligned to the onset of the videos (Figure 8A, 8C-D, 1-field: 44 / 198 = 22% of neurons, consistent with the main experiment results, cf. Figure 2), another set of neurons changed their firing pattern in accordance to the onset of each clip within the video (Figure 8B, 8D, 2-field: 10 / 198 = 5% of neurons, see summary table for temporal context cells and their overlap with TOJ cells in Table S2). These 2-field temporal context cells are able to more finely encode temporal information of video events in the presence of a clear event boundary within the video. This implies that the population of mPPC neurons adaptively encode for temporal structure and code the event boundaries of the finely segmented experiences dynamically, which might provide greater precision in temporal memory.

## DISCUSSION

In this study, we examined the role of mPPC neurons during the encoding and retrieval process of temporal order judgment. We found that mPPC neurons are strongly involved in both the encoding and retrieval of temporal information of events embedded in video material, with these neurons working in synchrony to enable successful judgment of temporal order. By identifying a subset of neurons that code for context within episodes, we found that neural coding for temporal context during experience may provide a scaffold for subsequent memory of event order. Below we discuss several implications of our discoveries.

We elucidated the neural activities for the memory traces during encoding and how they might be related to the activities upon retrieval. Inspired by the recent discovery of temporal context cells in the primate entorhinal cortex ^26^, we found a subset of mPPC neurons, labelled as temporal context cells, respond to the video context at the time of onset, but relaxing to baseline at different rates. This property has previously been shown in the entorhinal cortex ^26^ when still picture stimuli were used. The neurons we identified here are likely those that are triggered by the start of videos. Notably we conducted a control experiment in which we used 2-clip (i.e., across context) videos for encoding. As expected, we obtained evidence that some of the putative temporal context cells are 2-field and can code for the event boundaries of these segmented experiences. These temporal context cells, which are characterized by their wide spectrum of the past, are distinguished from neurons that are instantly responsive to hard or soft video boundaries in the human MTL ^35^ .With a proper set-up such as an ethogram^38,39^, it would be possible to expect certain cells are triggered by a certain feature and would relax back at different time scales ^27^.

We found that over two-thirds of the temporal context cells belonged to the class of TOJ cells (albeit a smaller proportion was found in experiment 2) and that the cells cover a wide spectrum of relaxation time. A population of temporal context cells with highly variable lengths of relaxation time could work as an instrumental encoding mechanism to process and encode the video experience as a coherent event ^40,41^. Since different neurons decay at a variety of rates, the animal could infer how far in the past certain frames were by processing which cells were active at which point. Howard and colleagues suggest such spectrum of time constants enable the brain to represent a Laplace transform of the organism’s past ^26,27^. This spectrum-based temporal record of the past has previously been observed in hippocampal neurons, which hold incremental timing signal for separate items within an event ^42^ as well as code for temporal-order relationships between events ^43,44^.

Another interpretation of the significance of such variable relaxation time could be related to the notion of temporal receptive windows ^45,46^. This phenomenon stipulates that cortical activities are organized in a hierarchical manner, with prefrontal and parietal areas exhibiting longer timescales ^46^. Converging with this idea, we trained a decoder to reveal that these neuron ensembles carry information about the passage of time embedded in the videos. We infer from the successful decoding of the second-by-second within-video temporal information that the mPPC neurons reflect information about the temporal unfolding of events across an extended window. This finding aligns with previous fMRI, single-unit, and electrocorticography evidence that cortical circuits can accumulate information over time following a hierarchical arrangement from early sensory areas with short processing timescales to higher-order areas such the mPFC and the mPPC hub with long processing timescales ^46,47^.

We sought to understand the mechanisms of mPPC involved in temporal memory retrieval by examining the single and ensemble neuronal activities. We found that a subpopulation of mPPC neurons which were identified as TOJ cells modulated their activity specifically when monkeys were judging the temporal order of probe frames. This neuronal activity was modulated by the correctness of response, implying the role of this brain region in memory retrieval. These findings align with fMRI studies identifying a similar function of mPPC during memory retrieval ^17,18^. Solely based on fMRI data, it is difficult to explore the underlying mechanism of how mPPC is involved in the reinstatement and recollection of temporal order memory ^48^. By contrast, we looked specifically into the temporal dynamics under TOJ memory recall and found that the level of neuronal synchrony increased gradually when temporal judgment was initiated. Irrespective of the memory outcome, the synchrony level of neuronal spikes peaked rapidly around 250 ms, indicating that decision evidence could be accumulated in a rapid manner, in line with effects reported in decision making ^49^, spatial ^50^ and working memory ^51^. Moreover, when we focused on the window leading to the moment of decision, we observed that correct decisions would trigger a significantly higher synchrony level of mPPC populations than incorrect trials immediately before the moment of decision. The effect is akin to the findings reported in the inferotemporal cortex where neuronal synchrony is enhanced for successful recognition of visual stimuli ^52^, implying that synchrony level of population determines the processes towards a correct decision ^53^. Additionally, we found that the neuronal synchrony level is higher for trials without a retention delay than for trials with a longer retention delay. These provided compelling evidence that the synchrony among the parietal neurons is an indicator of memory strength ^54^ of temporal order information. In the control analysis, we further ascertained that confounding effects from eye movements and scan paths do not account for the temporal order memory neuronal activity in a significant way.

To link it with the wider literature on temporal memory, we acknowledge a series of work that has indicated how cells in the hippocampus are recruited for encoding temporal information ^16,42,55^ and function as schema/event cells across extended time for experiences/memory ^56^. Multiple memory systems are also possible to support temporal order memory (e.g., with orbitofrontal cortex^57,58^ and contextual representation of events ^59^). One of the paths for future investigation should be focused on the interplay between MTL (e.g., time cells), medial parietal region ^60^, and the prefrontal region ^16^.

## STAR* METHODS

**KEY RESOURCES TABLE**

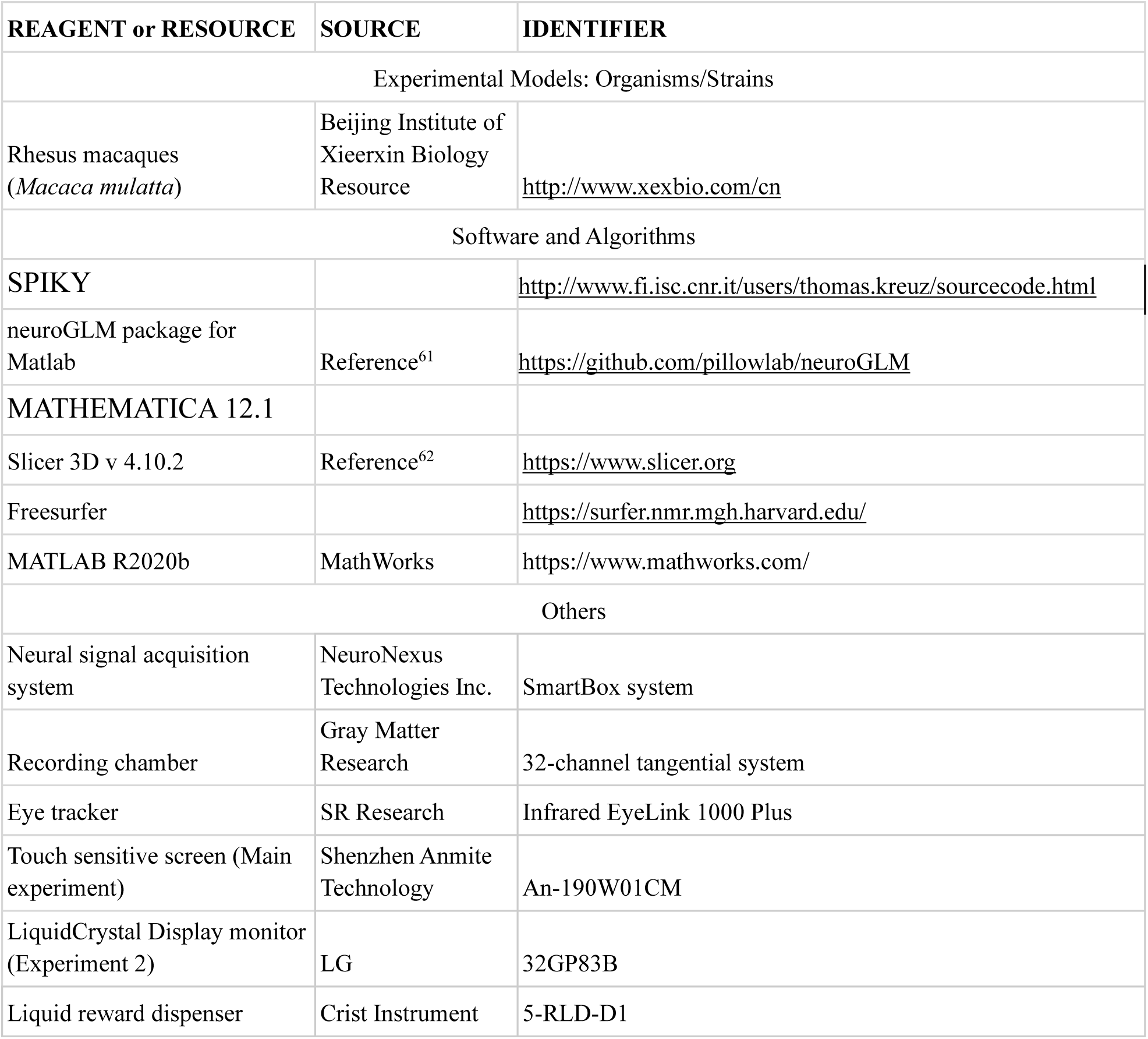

| REAGENT or RESOURCE | SOURCE | IDENTIFIER |
| --- | --- | --- |
| Experimental Models: Organisms/Strains |  |  |
| Rhesus macaques<br>(*Macaca mulatta*) | Beijing Institute of<br>Xieerxin Biology<br>Resource | http://www.xexbio.com/cn |
| Software and Algorithms |  |  |
| SPIKY |  | http://www.fi.isc.cnr.it/users/thomas.kreuz/sourcecode.html |
| neuroGLM package for<br>Matlab | Reference<sup>61</sup> | https://github.com/pillowlab/neuroGLM |
| MATHEMATICA 12.1 |  |  |
| Slicer 3D v 4.10.2 | Reference<sup>62</sup> | https://www.slicer.org |
| Freesurfer |  | https://surfer.nmr.mgh.harvard.edu/ |
| MATLAB R2020b | MathWorks | https://www.mathworks.com/ |
| Others |  |  |
| Neural signal acquisition<br>system | NeuroNexus<br>Technologies Inc. | SmartBox system |
| Recording chamber | Gray Matter<br>Research | 32-channel tangential system |
| Eye tracker | SR Research | Infrared EyeLink 1000 Plus |
| Touch sensitive screen (Main<br>experiment) | Shenzhen Anmite<br>Technology | An-190W01CM |
| LiquidCrystal Display monitor<br>(Experiment 2) | LG | 32GP83B |
| Liquid reward dispenser | Crist Instrument | 5-RLD-D1 |

## RESOURCE AVAILABILITY

**Lead contact.** Further information and requests for resources should be directed to and will be fulfilled by the lead contact, Sze Chai Kwok (<u>sze-chai.kwok@st-</u> hughs.oxon.org).

**Data and code availability.** Analysis code and processed data supporting the conclusions of this study are archived on https://github.com/.

## EXPERIMENTAL MODEL AND SUBJECT DETAILS

### Subjects

Three male rhesus macaques (*Macaca mulatta*) took part in this study. Two of them (5 and 5.5 years old, weighing 8.3kg and 8.6kg) participated in the main experiment and a third one (10 years old and 9.5kg) participated in experiment 2. The monkeys were housed in pairs but were singly housed during the period of this study. They were kept on a 12:12 (7:00am/7:00pm) light-dark circle and kept within the temperature range of 18 - 23℃ and humidity between 60% - 80%. The animals were fed twice a day with each portion of at least 180 g monkey chow and pieces of apple (8:30 am / 4:00 pm). Their water supply was only restricted during recording days. No food restriction was imposed throughout the study. The three monkeys were trained on a similar temporal order judgment task prior to this study ^28,63^. This study was conducted at the Nonhuman Primate Research Center at East China Normal University. All animal care, experimental, and surgical procedures were approved by the Institutional Animal Care and Use Committee (permission code: M020150902 & M020150902-2018) at East China Normal University.

### Experimental apparatus

For all recording sessions, the monkeys, head-refrained, sat in a custom-manufactured Plexiglas monkey chair (29.4 × 30.8 × 55 cm) inside a lead shielded chamber with ventilation during testing. In the main experiment, the monkeys responded with their right hand by touching the stimuli on a 19-inch touch-sensitive screen mounted on a stainless-steel platform. The monkeys were placed within their arm length to the touchscreen. In experiment 2, the monkey responded with its eye gaze by fixating for at least 1 s on a designated response area on a 31.5-inch LCD monitor (dimension = 714 × 428 mm; resolution = 1707 × 960). Monkeys’ eyes were about 60 and 62 cm away from the screen’s top edge and bottom edge. In both experiments, small amounts of water reward were delivered by a liquid dispenser according to a reward contingency described below.

### Surgical procedure

Anatomical T1 magnetic resonance images were acquired prior to the surgeries to guide us in the placement of the recording chambers. Detailed surgical procedures are reported in detail previously ^38^. In brief, the 32-channel recording chambers were placed over the medial posterior parietal cortex for monkey Jupiter (anteroposterior AP = - 16.4 mm, mediolateral ML = 5.8 mm; 28° to the right and 14° to the posterior of the transverse plane), monkey Mercury (- AP = - 15.4 mm, ML = 7.5mm; 25° to the right and 9.1° to the posterior), and monkey Mars (AP = - 16.9 mm, ML = -5.1 mm; 7° to the left and 48° to the posterior). Anatomical MRI images with a model of recording electrodes are shown in Figure 1B (upper panel). To confirm the electrode locations, we also acquired CT images (FOV = 8 cm, Voltage = 110kv, Current = 0.1mA) for each monkey and aligned the CT image with the MRI image to confirm our recorded sites at the end of the experiments (Figure 1B bottom for Jupiter and Mercury & 1C for Mars).

## METHOD DETAILS

### Temporal order judgment task

In the main experiment, two monkeys were trained to perform a temporal order judgment task based on naturalistic videos. In each trial, the monkey initiated a trial by pressing a colored rectangle in the center of the screen (0.3 ml water). Following a blank screen (range from 1s to 3s), an 8-s video was presented. After a short period of delay (no delay or 3.6s delay), two frames extracted from the video were displayed bilaterally on the screen for TOJ. The monkey was required to respond by touching the computer screen, and to choose the frame that was shown earlier in the video to get a reward (1.2 ml water), and the target frame remained alone for 4 s as positive feedback. If the monkey made an error, the screen would be blanked for 4 s with no reward. The temporal distances between the two frames were fixed in all trials (85 frames). To control for temporal similarity ^64^, we included two delayed conditions. In the immediate condition, the two probe frames were extracted from the first half of the video (i.e., 5th vs 90th frame), with zero retention delay between encoding and retrieval. In the delayed condition, the two probe frames were extracted from the second half of the video (i.e., 95th vs 180th frame), with a retention delay of 3.6 s.

In the first stage, we completed 24 sessions of Jupiter and 3 sessions of Mercury. 30 different videos were used in each block, and each video was shown once corresponding to two different conditions (i.e., immediate vs. delayed conditions). Each session contained 6 to 10 blocks (Jupiter: 8.417±1.100, Mercury: 7.333±1.528) depending on the monkey’s performance. In the second stage, we completed 18 sessions for both Jupiter and Mercury. 15 different videos were used twice with each video shown once for immediate and delayed conditions in each block. Each session contained 3 to 10 blocks (Jupiter: 8.444±1.723, Mercury: 5.667±1.138). In both stages, the same set of videos were randomized across different blocks and reused across three consecutive days. The difference between the stages was the number of videos used per session and the data from both stages was combined for analysis.

In experiment 2, an additional monkey was tested on the same paradigm. Each session contained 6 blocks, totaling 180 trials per day. The procedure was the same as the main experiment’s second stage except for two differences. Firstly, the monkey now needed to make their memory responses by fixating their gaze for at least 1s on one of two choice boxes (80 × 80 pixels) shown below the TOJ images. Secondly, the videos used in Experiment 2 consisted of two 4-s clips rather than one single 8-s video. The frames extracted for the temporal order judgment task followed the same rule as the main experiment and accordingly, all TOJ trials were performed across two contexts (across two clips).

### Eye movement recording

An infrared EyeLink 1000 Plus acquisition device (SR Research) was used to track eye positions at a sampling rate of 1,000 Hz. The illuminator module and the camera were positioned above the monkey’s head. An angled infrared mirror was used to capture and re-coordinate monkeys’ eye positions. The monkey’s right eye was tracked throughout the whole experiment session.

### Electrophysiology recording and spike sorting

On all three monkeys, we used 32-channel semi-chronic Microdrive systems (SC-32, GrayMatter Research) with 1.5 mm inter-electrode spacing for the recording. The headstage of the multichannel utility was connected to an acquisition system (SmartBox, NeuroNexus Technologies Inc., USA) via an amplifier Intan adapter (RHD2000, Intan Technologies) with 32 unipolar inputs. The microelectrode impedance of each channel was in the range of 0.5–2.5 MΩ and measured at the beginning of the session. Spike waveforms above a set threshold were identified with a 1,000 Hz online high-pass filter. Electrophysiological data collection was bandpass filtered from 0.1 to 5,500 Hz and digitized at 30 kHz.

## QUANTIFICATION AND STATISTICAL ANALYSIS

### Behavioral Data Analysis

Trials with reaction time longer than 10 s were excluded from analyses (1.71% in main experiment and 3.04% in experiment 2).

### Eye Movement Analysis

Raw eye movement data was converted from .edf format to .asc format. Saccades were identified by the EyeLink 1000 Plus acquisition system with a “SACC” marker during recording. The duration of a saccadic event was defined as the time elapsed starting from when the eye velocity exceeds 15° s^−1^ until when it slows down to below this velocity. We calculated the x and y coordinates for the eye position throughout the video and during the TOJ stage. Scan paths are then created by mapping the coordinates onto the video or TOJ images to produce eye position trajectories. The coordinates during blinks were filled with linear interpolation by using the coordinates 100 ms before and after a blink. For each trial, the time course of eye movement data was aligned with the neuronal data for the eye-spike analyses.

### Spike Detection and Sorting

The raw signal was filtered with a low-cut digital filter (4-pole Butterworth filter, 250 Hz), and we set 3 standard deviations as the threshold of detecting spikes. After removing high amplitude artifacts, we sorted spikes with the Standard E-M algorithm in Offline Sorter (Plexon). We used several criteria to quantify spike sorting quality and to select well-isolated single units, including signal-to-noise ratio, L-ratio, and isolation distance ^65^ (Figure S1). We treated clusters identified from the same electrodes on different days as separate neurons.

### Poisson GLM fitting and control analysis GLMs

We fit every single cell with Poisson GLM using *stepwiseglm* function in MATLAB. To exclude all the potential effects caused by behavior or other event period, we fit the model with following variables: block number within one session, reaction time, response correctness (correct vs incorrect), conditions (immediate vs delayed condition), touch side (left vs right), and several binary variables inputting the two states (yes/no) of pre-trial fixation, encoding period, delay period, TOJ period, feedback (reward vs punishment) and ITI period. Variables with an adjusted R^2^ smaller than 0.005 were excluded from the model, whereas variables with an adjusted R^2^ larger than 0.01were included. The firing rate of each event was calculated as the mean firing rate during that event. TOJ cells were defined as those cells who included TOJ period as a significant variable in the final model.

Moreover, using data from experiment 2, we performed a control analysis with GLMs on the TOJ firing rate including all the regressors in the original GLM (see above) but now also including several eye parameters such as saccadic frequency and length of scan paths measured. The purpose of this control GLM was to leverage eye movement data to regress out any confounding role of eye movement parameters. The exact parameters included in the model were: frequency of saccades, frequency of fixation, total length of saccade path (in pixel), and total fixation duration (in second) during the TOJ period. Similar to the main GLM model, variables with an adjusted R2 smaller than 0.005 were excluded from the model, whereas variables with an adjusted R2 larger than 0.01were included. In this control analysis, TOJ cells were more strictly defined as those cells who included TOJ period as a significant variable without having any of the eye parameters included as a significant variable.

### Mahalanobis Distance Index

We first calculated the mean firing rate of each neuron recorded in that session for Video watching and TOJ period trial by trial. Next, the squared Mahalanobis distance of each trial was calculated with the MATLAB function ‘*mahal*’. Since the number of neurons are different from session to session, we divided the Mahalanobis distance with two times of cell number in that session for comparison across sessions.

### SPIKE-distance

The SPIKE-distance measures the relative spike timing between spike trains normalized to local firing rates. For each spike of each pair of spike trains recorded in the same session, we calculated the time difference between the nearest following spikes of two spike trains and the time difference between the nearest preceding spikes of two spike trains. These distances are interpolated between spikes using all time differences to the preceding spike and the following spike. The instantaneous weighted spike time difference for a spike train can be determined via the interpolation from one difference to the next. For each session, we averaged over all pairs of neurons to acquire the averaged SPIKE-distance value for each time point. The pairwise SPIKE-distance profile is subsequently obtained by averaging the weighted spike time differences, normalized to the local firing rate average, and weighted each profile by the instantaneous firing rates of the two spike trains. Averaging over all pairwise SPIKE- profiles results in a multivariate SPIKE-profile. We obtained the SPIKE-distance value for each pair of two spike trains and represented all pairwise averaged dissimilarity profiles in a n x n matrix (n denotes the number of neurons recorded in one session). Calculation of SPIKE-distance and pairwise SPIKE-distance for all neurons in the same session was done with SPIKY ^66^. For interpretation, the smaller the value is, the more similar the spike trains are, and thus there is a higher synchrony between spike trains.

### Temporal Context Cell Fitting

We analyzed the spike data of the monkeys via a maximum likelihood estimation script run in MATHEMATICA 12.1. In order to determine whether a neuron had a time-locked response to the onset of video clips, we calculated model fits of nested models for each neuron across all trials considering the time from the onset of image presentation to 8s after image presentation. The nested models contain three models.

1. The constant model, 𝐹_𝑐𝑜𝑛𝑠𝑡_(𝑡; 𝑎_0_) = 𝑎_0_ , which is only determined by a single parameter 𝑎_0_ that predicted the average firing rate at each time 𝑡.
2. The Gaussian model,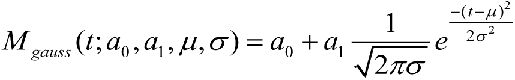
3. The ex-Gaussian model, 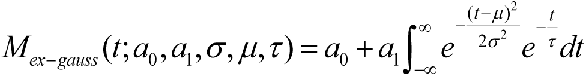

which is represented by the convolution of a Gaussian function with an exponentially decaying function.

Since the ex-Gaussian model degrade into either a Gaussian function (as *τ* →0) or an exponential function starting at *µ* (as *σ* → 0), this model performs well in describing both temporal context cells and time cells. As such, neurons which are better fitted by the ex-Gaussian model are considered responsive. We selected neurons with three criteria: 1) were better fitted by the ex-Gaussian model at the 0.05 level, 2) changed their firing rate by at least 2 Hz, 3) reached a firing rate of at least 4 Hz. Fits of nested models for each neuron are analyzed via a likelihood ratio test.

Regarding Experiment 2, we also studied whether neurons have a time-locked response to the onset of each 4-s clip. To account for the two-field responsive property, we extended our analysis by incorporating a two-field ex-Gaussian model into our nested model comparison:

1. 4) The two-field ex-Gaussian model,

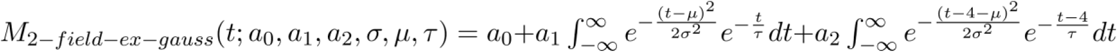

This model introduces a single additional parameter, compared to the one-field model. We identified responsive neurons by applying the same selection criteria used for the one-field analysis but now selecting neurons that showed superior fit to the two-field ex-Gaussian model.

### Linear Discriminant Analysis

To decode temporal information from the population activity of mPPC neurons, we followed the analysis method of the previous study ^26^. We trained the classifier with even trials and tested the temporal information in odd trials. For each 8 s video in each trial, we took 250 ms as the time bin and divided the whole encoding period into 32 bins. For each bin of each trial, the firing rate of all neurons in that session was calculated. One element of testing data or training data is the average firing rate of each time bin for each trial across all neurons. Due to the number of trials that range from session to session, we randomly select 200 trials when the number of trials in one session is larger than 200. While the number of trials is fewer than 200, we bootstrapped it to 200 trials. In this case, some trials would have been selected twice. LDA was implemented using the MATLAB function “classify”, and we used linear estimation for our dataset. To assess the actual error in each classification, we calculated the absolute temporal distance between the actual time and decoded time in each time bin and got the averaged value across all the time bins. Besides, due to the error in the first period of the video being lower than the rest of the video for two monkeys (Figure S3), we repeat the same analysis when removing the time bins from the earlier part of the video at the step of 0.5 s. To quantify the success of LDA, we permuted time labels of training data and repeated the analysis 1000 times. The classifier’s performance was considered significantly better than chance if fewer than 10 of 1000 permutations gave a better result than the real data.

## SUPPLEMENTAL INFORMATION

Supplemental information can be found online.

## ACKNOWLEDGMENTS

This work received support from the National Natural Science Foundation of China (32071060), Jiangsu Provincial Department of Science and Technology (BK20221267), and internal funding from School of Psychology and Cognitive Science at East China Normal University (SCK). SZ received support from the Kobayashi Foundation. We would like to extend our heartfelt gratitude to Professor Edmund Rolls for passing on his NHP electrophysiology mastership to the team. We would like to dedicate this work to the late Professor Yong-di Zhou, who founded the Nonhuman Primate Research Center at East China Normal University.

## AUTHOR CONTRIBUTIONS

Conceptualization, SZ, SCK; methodology, SZ, ZJ, XZ, WL, NS, JL, TJM, MK, SCK; investigation, SZ, ZJ, XZ, NS, JL, MK; formal analysis, SZ, CW; visualization, SZ, CW; writing – original draft, SZ, CW, SCK; writing – review & editing, SZ, SCK; supervision, SCK; funding acquisition, SCK.

## DECLARATION OF INTERESTS

The authors declare no competing interests.

## Notes

### Competing Interest Statement

The authors have declared no competing interest.

### Summary of Updates

We have trained a new monkey on a variant of the same memory task that relies on eye-movement rather than manual responses. We then operated on this monkey with the same electrophysiology recording set-up. We then obtained 198 new neurons from this monkey, while we simultaneously monitored the animals eye position and saccadic scan paths over 21 experimental sessions. This new data helps us ascertain how eye positions and oculomotor behavior might be driving the neuronal activities during encoding and memory retrieval. Specifically, we analyzed the animals task-related eye position and scan path trajectory, as well as the new neurons corresponding firing pattern. We presented the firing rate as a function of either fixation or scan path, separately for correct and incorrect responses (see below for example cells, or our new Figure 7). We also included a movie for illustration of a complete trial with eye movement tracking (Movie S1). From this new data, we can see that the animal overtly attended to the frames while they were performing the memory task.

