## Supplement file for "Neural signatures for temporal order memory in the macaque medial posterior parietal cortex"

This manuscript contains two (2) supplemental figures, two (2) supplemental tables, and one (1) supplemental movie.

Supplemental Figure S1:
Spike sorting metrics. Is associated with main Figure. 1.

Supplemental Figure S2:
Decoding errors remain significantly lower than permutated data with different numbers of bins discarded. Is associated with main Fig. 5.

Supplemental Table S1:
Summary of temporal order judgment (TOJ) cells identified by main GLM and control GLM with eye movement parameters.

Supplemental Table S2:
Summary of proportion of temporal context (TC) cells overlapped with TOJ cells for the 3 monkeys.

Supplemental Movie S1:
A movie displaying a complete example trial with eye movement tracking.

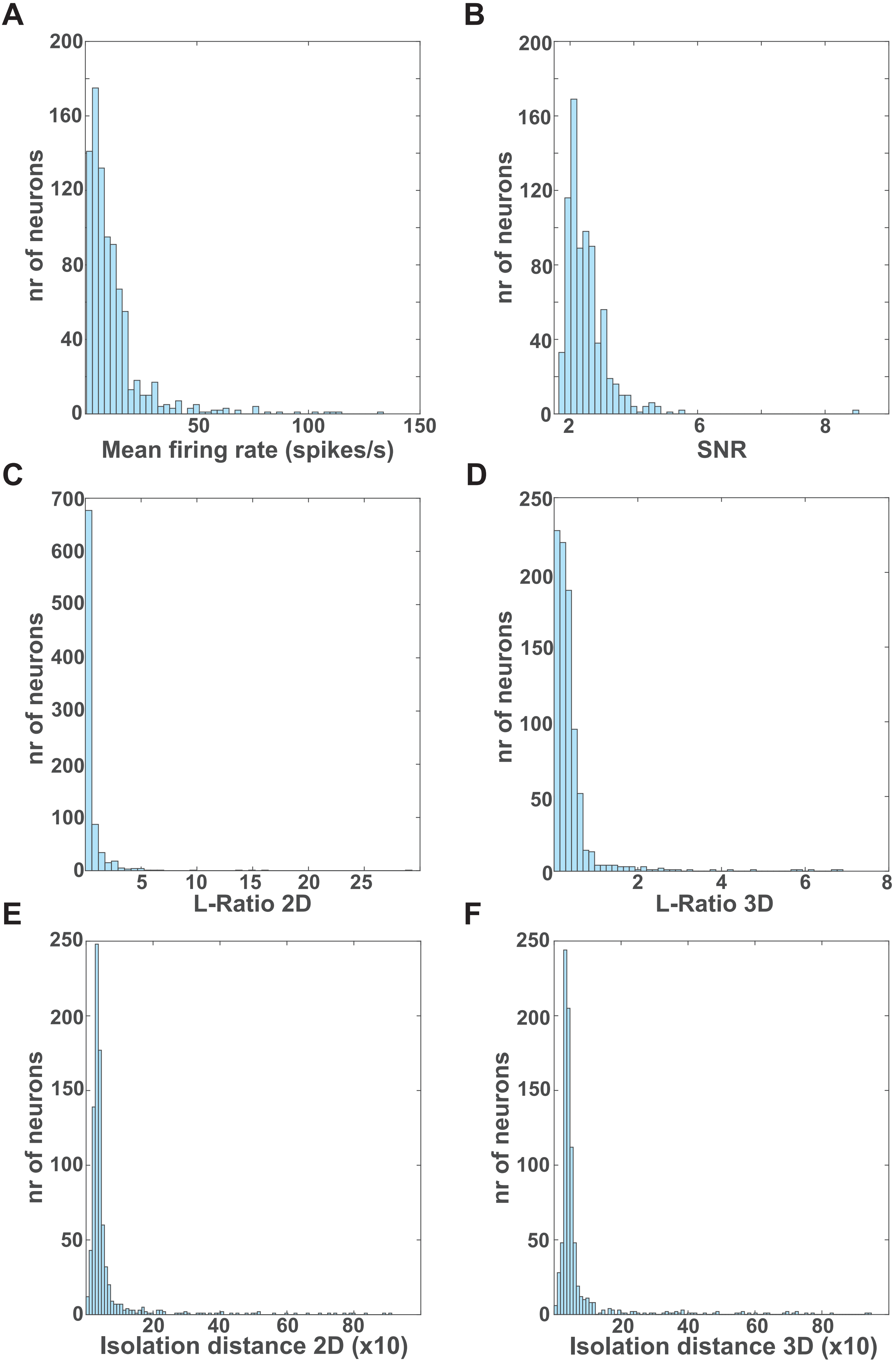

A
nr of neurons
Mean firing rate (spikes/s)
B
nr of neurons
SNR
C
nr of neurons
L-Ratio 2D
D
nr of neurons
L-Ratio 3D
E
nr of neurons
Isolation distance 2D (x10)
F
nr of neurons
Isolation distance 3D (x10)

**Supplemental Figure S1 Spike sorting metrics.**

(A) Mean firing rate of all recorded neurons from 3 monkeys (401 and 275 cells from Main experiment and 198 cells from Experiment 2). The average firing rate with SD was 12.29±14.38 (spikes/s). (B) Histogram showing numbers of neurons as a function of signal-to-noise ratio (SNR) of the mean waveform of neurons. The mean SNR with SD was 2.50±0.73. (C) Histogram showing numbers of neurons as a function of L-Ratio value calculated based on 2D clusters. The average value with SD was 0.59±1.56. (D) Histogram showing numbers of neurons as a function of L-Ratio value calculated based on 3D clusters. The average value with SD was 0.39±0.63. (E) Histogram showing numbers of neurons as a function of isolation distance calculated based on 2D clusters. The average value with SD was 6.20±10.76. (F) Isolation distance calculated based on 3D clusters for all neurons. The average value with SD was 7.12±11.49, indicating that clusters associated with single neurons were well separated from all other clusters.

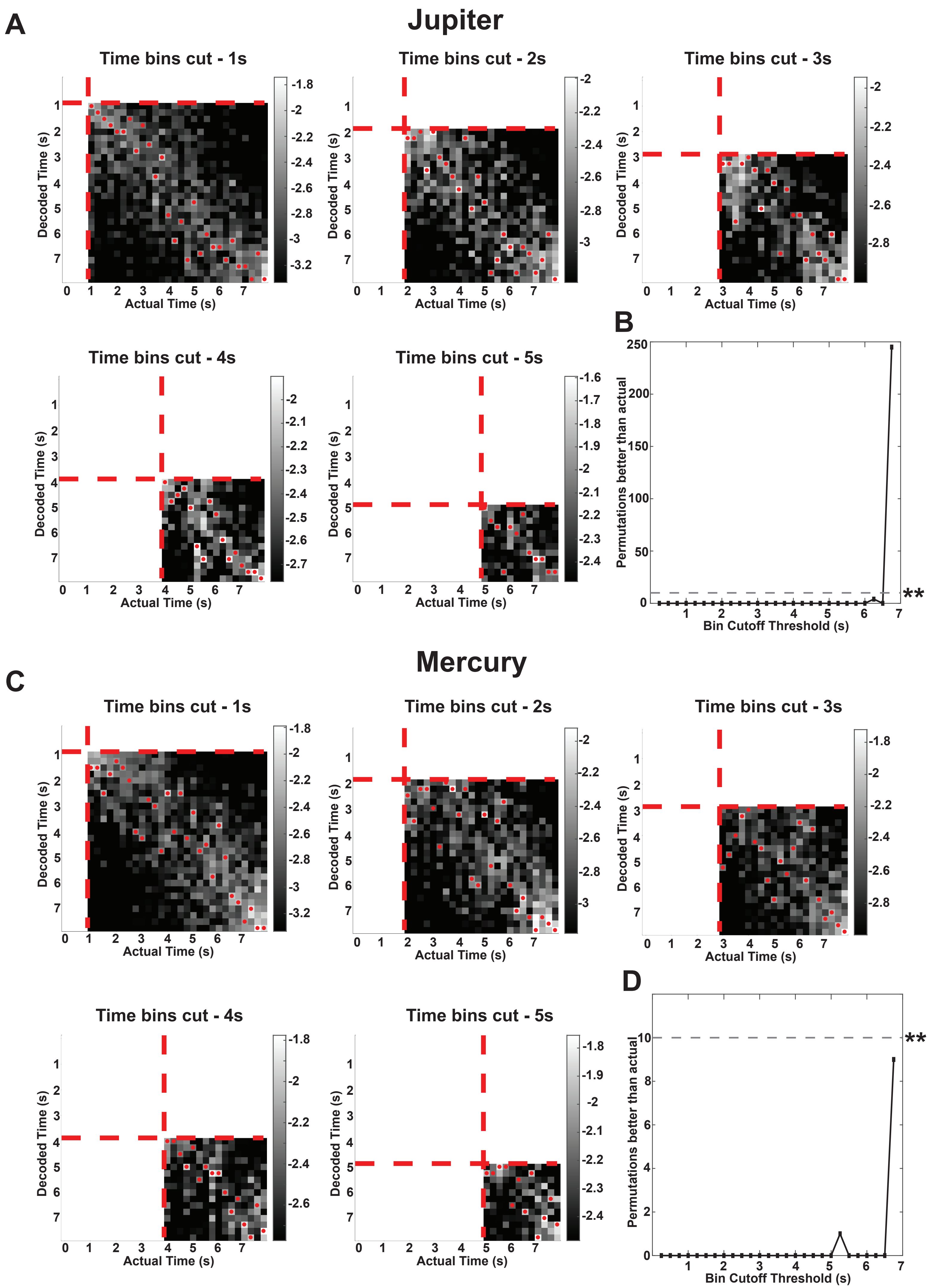
A
Jupiter
Time bins cut - 1s
Time bins cut - 2s
Time bins cut - 3s
Time bins cut - 4s
Time bins cut - 5s
Decoded Time (s)
Actual Time (s)
B
Permutations better than actual
Bin Cutoff Threshold (s)
**
C
Mercury
D

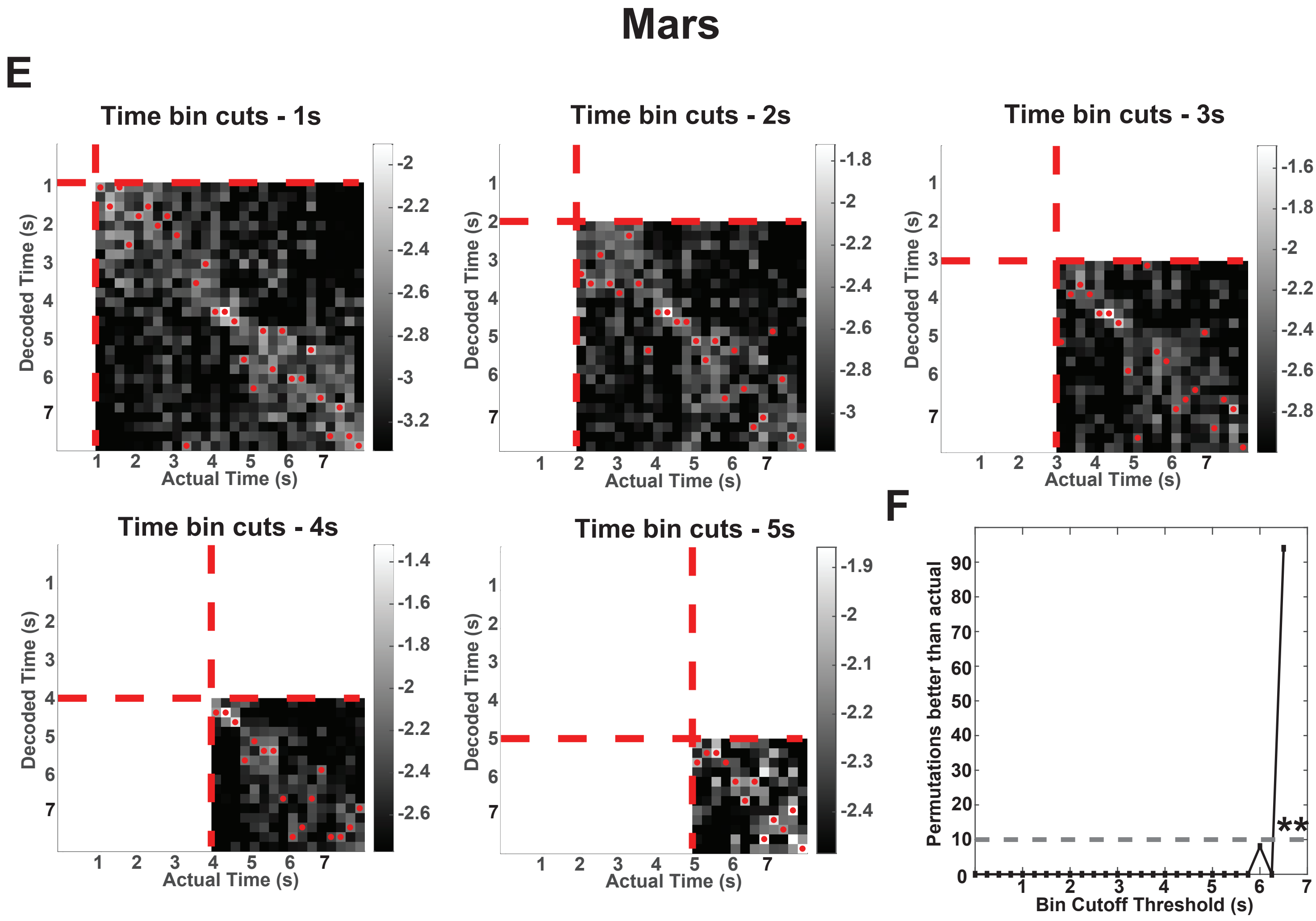
Mars
E
Time bin cuts - 1s
Time bin cuts - 2s
Time bin cuts - 3s
Time bin cuts - 4s
Time bin cuts - 5s
Decoded Time (s)
Actual Time (s)
F
Permutations better than actual
Bin Cutoff Threshold (s)
**

**Supplemental Figure S2 Decoding error remain significantly lower than permutated data with different numbers of bins discarded (A: Jupiter, C: Mercury, E: Mars).**

The posterior probability as bins dropped (A: Jupiter, C: Mercury, E: Mars) are shown. For all three monkeys, the top left panel refers to decoded posterior probability with the first 1s data cut off, with the other panels corresponding to decoded posterior probability with the first 2s, 3s, 4s, and 5s data cut off. B, D and F show their corresponding decoder performance (B: Jupiter, D: Mercury, F: Mars). The decoded time with the highest probability in each time bin is marked with a red dot. Each second was marked with red arrows along the axis. All decoder performance were significantly better than chance even when we excluded up to the first 6s. The line marked by '**' is significance at $p < 0.01$.

**Table S1. Summary of temporal order judgment (TOJ) cells identified by main GLM and control GLM for eye movement parameters.**

| | | | | TOJ cells result based on Poisson GLMs | | | |
|---|---|---|---|---|---|---|---|
| | | | | Original GLM | | GLM with eye parameters | |
| **Monkey** | **Experiment** | **Eye data availability** | **Total cells recorded** | **# TOJ cells** | **% TOJ cells** | **#TOJ cells** | **% TOJ cells** |
| Jupiter | Main | No | 401 | 278 | 69.3% | N/A | N/A |
| Mercury | Main | No | 275 | 169 | 61.5% | N/A | N/A |
| Mars | Exp2 (2-segment videos) | Yes | 198 | 58 | 29.3% | 41 | 20.7% |
| **Total** | | | 874 | 505 | | | |

**Table S2. Summary of proportion of temporal context (TC) cells and their overlap with TOJ cells for the 3 monkeys.** For Mars, the 54 TC cells consisted of 44 2-field cells and 10 1-field cells.

| | | | | Temporal context (TC) cell | | |
|---|---|---|---|---|---|---|
| **Monkey** | **Experiment** | **Total cells recorded** | **# TC cells** | **% of TC / Total cells** | **(TOJ cells ∩ TC cells) / TOJ cells** | **(TOJ cells ∩ TC cells) / TC cells** |
| Jupiter | Main | 401 | 66 | 16.5% | 15.5% | 65.2% |
| Mercury | Main | 275 | 45 | 16.4% | 20.7% | 77.8% |
| Mars | Exp2 (2-segment videos) | 198 | 54 | 27.3% | 22.4% | 24.1% |
| **Total** | | 874 | 165 | | | |

**Supplemental Movie S1:**

A movie displaying a complete example trial with eye movement tracking. The trial events included fixation, video (encoding), delay (0-s delay), temporal order judgement (two concurrently presented frames), and feedback. The green moving dot is the monkey's eye gaze (right eye).
